# Molecular Detection and Antibiogram Assay of Escherichia coli Isolated from Poultry Farm Environments

**DOI:** 10.64898/2026.08.05.743159

**Authors:** Anna Purnna Ray, Md. Rimon Bhuiyan, Satyajit Roy, Hridoy Roy, Md. Sabbir Hossain, Md. Rashedul Kabir Mondol, K.M. Mozaffor Hossain

## Abstract

**Background:** Escherichia coli is a common environmental and zoonotic bacterium that serves as an important indicator of fecal contamination and antimicrobial resistance (AMR) in poultry production systems. The emergence of multidrug-resistant (MDR) E. coli in poultry farm environments poses a significant One Health threat to animal, human, and environmental health.

**Objective:** This study aimed to isolate, identify, determine the prevalence, assess the antibiotic susceptibility pattern, and molecularly confirm Escherichia coli isolated from poultry farm environments in Lalmonirhat district, Bangladesh.

**Materials and Methods:** A cross-sectional study was conducted from January to June 2024 using 60 environmental samples comprising water (n = 20), soil (n = 20), and bird-dropping (n = 20) samples collected from poultry farms in five upazilas of Lalmonirhat district. Isolation and identification of E. coli were performed using standard cultural, morphological, and biochemical techniques. Antimicrobial susceptibility was determined by the Kirby–Bauer disc diffusion method following CLSI guidelines. Ten randomly selected isolates were confirmed by polymerase chain reaction (PCR) targeting the species-specific uidA gene.

**Results:** The overall prevalence of E. coli was 50% (30/60). Source-wise prevalence was highest in bird-dropping samples (75%), followed by water (45%) and soil (30%). Area-wise prevalence ranged from 33.33% in Patgram to 66.67% in Hatibandha. The isolates exhibited the highest resistance to oxytetracycline (66.67%) and ciprofloxacin (60%), while the highest susceptibility was observed to erythromycin (63.33%), neomycin (56.67%), and amoxicillin (56.67%). All ten isolates subjected to PCR produced the expected 486 bp uidA gene amplicon, confirming their identity as E. coli.

**Conclusion:** The findings demonstrate a considerable prevalence of antimicrobial-resistant E. coli in poultry farm environments of Lalmonirhat district. Strengthening farm biosecurity, improving hygiene and waste management, implementing antimicrobial stewardship, and maintaining continuous molecular surveillance are essential to minimize the dissemination of resistant E. coli within a One Health framework.

## 1. INTRODUCTION

Bangladesh is predominantly an agrarian country, and its poultry sector has emerged as one of the fastest-growing components of national agriculture, expanding at an estimated 15% annually and contributing approximately 1.6% to the national gross domestic product (Department of Livestock Services, 2020–2021). The sector provides high-quality, digestible protein and essential micronutrients such as vitamins A and B12, iron, zinc and iodine, and chicken remains the most widely consumed meat product both domestically and globally (Adams and Moss, 2008). The country presently maintains around 90,000 registered poultry farms in addition to numerous informal operations, with a total poultry population of roughly 356.318 million birds comprising broilers, layers and other categories (DLS, 2020–2021). According to the DLS Livestock Economy Report, Bangladesh consumes about 3,340 tonnes of poultry meat daily, amounting to approximately 1.26 million tonnes annually, with poultry contributing 22–27% of the country’s animal protein supply and around 37% of total meat production (DLS, 2020–2021).

Despite its economic and nutritional importance, the poultry industry is also a significant reservoir of zoonotic and antibiotic-resistant bacteria, chief among them Escherichia coli (Kaper, Nataro and Mobley, 2004). E. coli is a normal commensal of the gastrointestinal tract of humans, poultry and other warm-blooded animals, but certain strains are capable of causing severe extra-intestinal and intestinal disease, including septicaemia, meningitis, endocarditis, urinary tract infection and epidemic diarrhoea in humans (Kaper, Nataro and Mobley, 2004), and yolk sac infection, omphalitis, respiratory tract infection, septicaemia, polyserositis, enteritis, cellulitis and salpingitis in birds (Rosario et al., 2004). Among avian strains, avian pathogenic E. coli (APEC), a type of extra-intestinal pathogenic E. coli (ExPEC), causes colibacillosis, typically presenting as acute fatal septicaemia or subacute fibrinous pericarditis, airsacculitis, salpingitis and peritonitis in broiler chickens aged four to six weeks (Alexander et al., 2017). The continued rise of multidrug-resistant E. coli strains has been linked to a marked increase in morbidity and mortality in humans, in part because resistance genes can be disseminated between farm-animal and human-associated strains through specific plasmid lineages (Been et al., 2014).

The growing trend of antimicrobial resistance (AMR) is driven substantially by the indiscriminate and excessive use of antibiotics in livestock and poultry production, both for disease prevention and as growth promoters (Economou and Gousia, 2015). Commonly abused agents include oxytetracycline, ciprofloxacin, neomycin, gentamicin, enrofloxacin, doxycycline, colistin, streptomycin, tylosin, nitrofurans and chloramphenicol (Habib et al., 2021). Several interlinked practices contribute to this problem, including unnecessary antibiotic prescriptions, extensive use in aquaculture and livestock, incomplete treatment courses, poor hygiene, bacterial mutation and a diminishing pipeline of new antibiotic discoveries (World Health Organization, 2020). Because inappropriate or excessive antibiotic use is a principal driver of resistant-bacteria emergence, antibiotic resistance management has become a central pillar of contemporary public-health strategy (Read and Woods, 2014).

E. coli naturally inhabits the intestines of warm-blooded animals and is therefore continually exposed to antimicrobial agents administered to the host; this sustained selective pressure favours the emergence and persistence of resistance (Levy and Ruiz, 2014; Dahal, 2017). Antibiotic-resistant E. coli can also transfer resistance determinants to other bacterial species through horizontal gene transfer, and natural environments such as soil, water and wastewater-treatment systems have been described as "genetic reactors" in which such exchanges occur frequently, analogous to processes within the intestinal tract (Tenaillon et al., 2010; Sterblad et al., 2000). Pathogenic and resistant E. coli enter the environment through manure, animal waste, slaughterhouse effluents and wastewater discharges (Balière et al., 2015), and although much research attention has focused on the clinical dimensions of E. coli infection, comparatively little has examined its prevalence and persistence in environmental settings such as farm soil, water bodies and litter (Kaper et al., 2004; Jang et al., 2017). This gap is particularly concerning given that antimicrobial resistance is a multifaceted One Health problem interlinking human health, animal health, food safety and the environment, and given that the continuous release of antibiotics and antibiotic residues into wastewater may further compromise environmental bacterial populations and freshwater ecosystems (Roose-Amsaleg and Laverman, 2015; Dargatz et al., 2000).

In many Asian countries, live poultry markets and farms represent a critical link in the poultry supply chain, and overcrowded, unhygienic holding conditions promote the survival and spread of environmental bacteria such as E. coli (Shao et al., 2016). Direct contact between farm workers, consumers and live or freshly slaughtered birds further increases the likelihood of transmitting antibiotic-resistant bacteria to humans through handling or consumption (Sackey et al., 2001). In Bangladesh, as in many other low- and middle-income countries, AMR is emerging as a critical public-health issue, yet awareness of bacterial contamination and infection risk on poultry farms remains limited (Hossain et al., 2020). Polymerase chain reaction (PCR) has become a widely applied molecular tool for bacterial detection because it enables rapid, sensitive and specific amplification of species-specific genes, offering an important complement to conventional culture-based identification (Jang et al., 2017).

Against this background, the present study was undertaken to investigate the prevalence, antibiotic-resistance profile and molecular identity of Escherichia coli isolated from soil, water and bird-dropping samples collected from poultry farms in Lalmonirhat district, Bangladesh, an area in which such baseline data have not previously been reported (Ibrahim et al., 2023). The specific objectives were to: (i) isolate and identify E. coli from the poultry farm environment (Cheesbrough, 2006); (ii) determine the antimicrobial-resistance pattern of the isolates by the disc diffusion method (CLSI, 2021); and (iii) characterise the isolates molecularly by PCR targeting the uidA gene (Feng, Weagant and Jinneman, 2020).

## 2. MATERIALS AND METHODS

### 2.1 Study area, duration and sample collection

This cross-sectional study was carried out between January and June 2024 in the Microbiology Laboratory, Department of Veterinary and Animal Sciences, University of Rajshahi. A total of 60 environmental samples, comprising 20 water samples, 20 soil samples and 20 bird-dropping samples, were randomly collected from poultry farms (broiler, layer and Sonali/coloured-bird farms) located in five upazilas of Lalmonirhat district, namely Lalmonirhat Sadar, Aditmari, Kaliganj, Hatibandha and Patgram, with 12 samples (4 water, 4 soil, 4 bird droppings) obtained from each upazila. Samples were collected aseptically using sterile plastic bags, Eppendorf tubes and cotton swabs and transported to the laboratory in ice boxes for immediate processing.

### 2.2 Sample preparation and culture media

A sterile cotton swab moistened with sterile normal saline was used to swab each sample, which was then immersed in 10 ml of normal saline and subsequently enriched in 60 ml of buffered peptone water, followed by incubation at 37°C for 18 hours. Standard bacteriological media, including nutrient broth, nutrient agar, MacConkey agar, Eosin Methylene Blue (EMB) agar, Triple Sugar Iron (TSI) agar, Simmons citrate agar, brilliant green agar and Mueller–Hinton agar, were prepared according to the manufacturers’ instructions and sterilised by autoclaving at 121°C and 15 psi for 15 minutes (Cheesbrough, 2006).

### 2.3 Isolation, cultural and morphological identification

A loopful of the enriched culture was streaked onto EMB agar and incubated aerobically at 37°C for 18–24 hours (Cheesbrough, 2006). Colonies presumptively identified as E. coli, appearing dark with a characteristic metallic green sheen, were sub-cultured to obtain pure isolates (Bell and Kyriakides, 2009). Colony morphology (size, shape, texture, elevation, margin, colour and opacity) was recorded after 24 hours of incubation on nutrient agar, MacConkey agar and brilliant green agar (MacFaddin, 2000). Gram’s staining was performed according to standard procedure (Cheesbrough, 2006) to determine cell morphology and Gram reaction, and motility was assessed by the hanging-drop technique using a compound microscope at 100× magnification with immersion oil (MacFaddin, 2000).

### 2.4 Biochemical characterisation

Isolates were subjected to a battery of biochemical tests, including the catalase test, indole test, methyl red (MR) test, Voges–Proskauer (VP) test, Simmons citrate utilisation test, and TSI agar slant reaction, together with fermentation of five basic sugars (dextrose, lactose, sucrose, maltose and mannitol) using Durham’s tubes to detect acid and gas production (MacFaddin, 2000). Reactions were interpreted according to standard colour-change and gas-production criteria (Cheesbrough, 2006).

### 2.5 Antibiogram assay

Antibiotic susceptibility of the confirmed isolates was determined by the Kirby–Bauer disc diffusion method on Mueller–Hinton agar (Bauer et al., 1966), in accordance with the Clinical and Laboratory Standards Institute guidelines (CLSI, 2021). Seven commercially available antibiotic discs (Hi-Media, India)- ciprofloxacin (5 µg), amoxicillin (30 µg), doxycycline (30 µg), oxytetracycline (30 µg), levofloxacin (5 µg), neomycin (30 µg) and erythromycin (15 µg)-representing antimicrobial classes commonly used in poultry and veterinary practice were applied to inoculated plates (Habib et al., 2021). Plates were incubated at 37°C for 16–18 hours, after which the diameter of the zone of inhibition was measured to the nearest millimetre and isolates were classified as resistant, intermediate or sensitive according to the CLSI interpretive breakpoints summarised in Table 1 (CLSI, 2021).

**Table 1.** Zone-diameter interpretive standards for antimicrobial susceptibility testing.

| Antimicrobial agent | Symbol | Disc content (µg) | Resistant (mm) | Intermediate (mm) | Sensitive (mm) |
| --- | --- | --- | --- | --- | --- |
| Ciprofloxacin | CIP | 5 | ≤15 | 16–20 | ≥21 |
| Amoxicillin | AMX | 30 | ≤13 | 14–17 | ≥18 |
| Doxycycline | DO | 30 | ≤10 | 11–13 | ≥14 |
| Oxytetracycline | TE | 30 | ≤11 | 12–14 | ≥15 |
| Levofloxacin | LVX | 5 | ≤13 | 14–16 | ≥17 |
| Neomycin | N | 30 | ≤12 | 13–16 | ≥17 |
| Erythromycin | E | 15 | ≤13 | 14–22 | ≥23 |

### 2.6 Stock culture maintenance

Confirmed isolates were preserved in 50% sterile buffered glycerine (equal parts glycerine and phosphate-buffered saline) at −20°C, a method capable of maintaining bacterial viability and phenotypic stability for six months to one year (Hossain et al., 2021).

### 2.7 Molecular detection by polymerase chain reaction (PCR)

Of the 30 culturally, morphologically and biochemically confirmed E. coli isolates, 10 were randomly selected for molecular confirmation (Jang et al., 2017). Genomic DNA was extracted by the boiling method: a pure colony of each isolate was grown in nutrient broth at 37°C for 8 hours, after which 1 ml of the culture was centrifuged at 10,000 rpm for 5 minutes; the pellet was washed with distilled water, resuspended in 200 µl of distilled water, boiled for 10 minutes, immediately chilled on ice for 10 minutes and centrifuged again, and the resulting supernatant was used as the DNA template (Hossain et al., 2021).

PCR amplification targeted the species-specific uidA gene (Feng, Weagant and Jinneman, 2020) using the forward primer 5′-ATCACCGTGGTGACGCATGTCG-3′ and the reverse primer 5′-CACCACGATGCCATGTTCATCTGC-3′, yielding an expected amplicon of 486 base pairs (bp) (Table 2) (Hossain et al., 2021). Each 12.5 µl reaction contained 6.25 µl of 2× PCR master mix (Promega, USA), 0.5 µl each of forward and reverse primer, 2.0 µl of DNA template and 3.25 µl of nuclease-free water (Sambrook and Russell, 2001). Amplification was performed in a thermal cycler (ASTEC, Japan; Model 482) under the cycling conditions summarised in Table 3 (Hossain et al., 2021). Amplified products were resolved by electrophoresis on a 1.5% agarose gel alongside a 100 bp DNA ladder (Promega, USA), stained with ethidium bromide (0.5 µg/ml) and visualised under a UV transilluminator (Bio-Rad, USA) (Sambrook and Russell, 2001); isolates producing the expected 486 bp band were considered positive for the uidA gene and thus confirmed as E. coli (Jang et al., 2017).

**Table 2.**
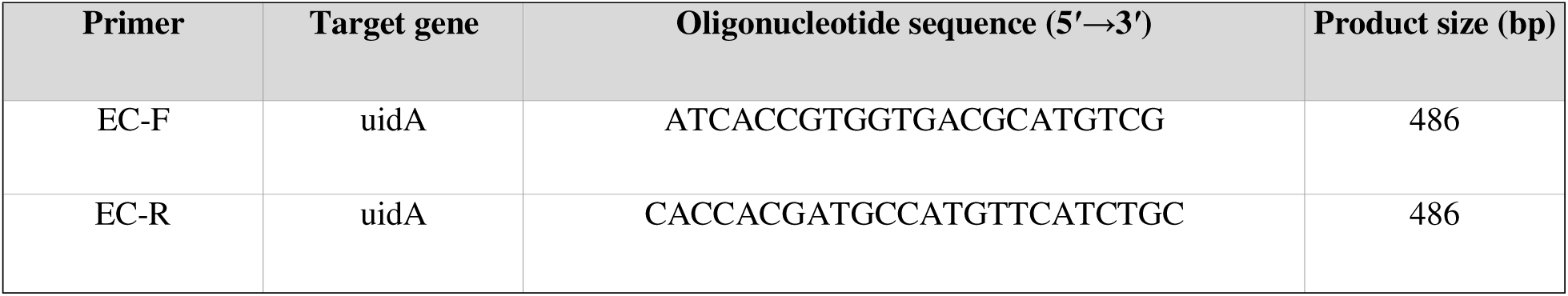
Primer sequences used for PCR identification of E. coli.

**Table 3.** PCR cycling conditions for amplification of the uidA gene.

| Step | Temperature (°C) | Time (min) | No. of cycles |
| --- | --- | --- | --- |
| Initial denaturation | 95 | 7 | 1 |
| Denaturation | 94 | 1 |  |
| Annealing | 55 | 1 | 35 |
| Extension | 72 | 1 |  |
| Final extension | 72 | 7 | 1 |
| Holding | 4 | ∞ | — |

### 2.8 Statistical treatment

Data on prevalence and antibiotic susceptibility were expressed as percentages, calculated as the number of positive samples or isolates divided by the total number tested, multiplied by 100 (Thrusfield, 2013).

## 3. RESULTS

### 3.1 Cultural and morphological characteristics

Growth of E. coli in nutrient broth was indicated by diffuse turbidity, occasionally accompanied by pellicle formation. On nutrient agar, colonies appeared smooth, circular and greyish-white; on EMB agar, colonies were smooth, circular and greenish-black with a characteristic metallic sheen; on MacConkey agar, colonies were bright pink owing to lactose fermentation; and on brilliant green agar, colonies were yellow, with the medium changing from red to yellow. Gram’s staining revealed Gram-negative, small rod-shaped cells arranged singly or in pairs, and the hanging-drop motility test confirmed that the isolates were motile.

**Figure 1.**
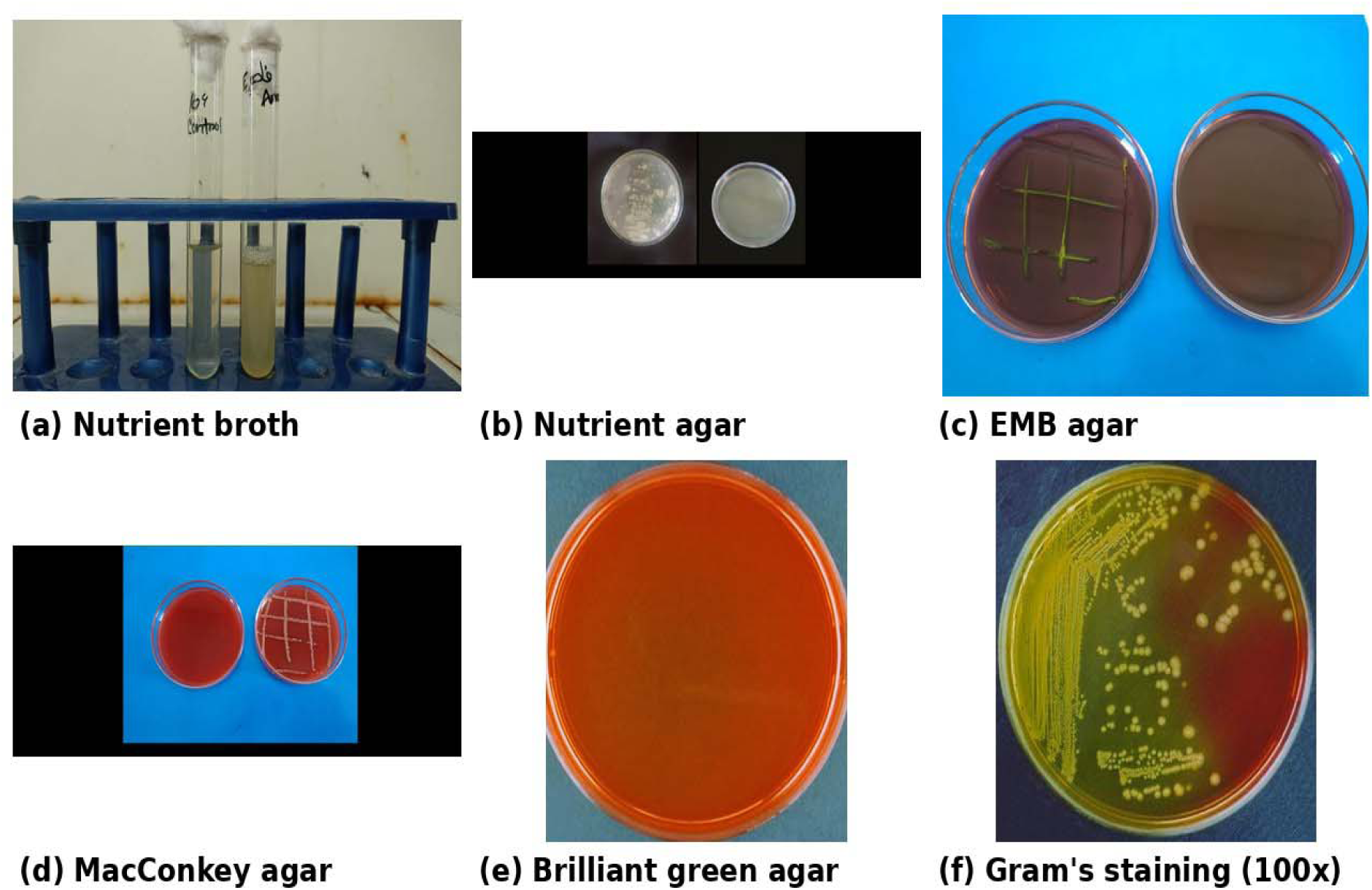
Cultural and morphological characteristics of E. coli isolated from the poultry farm environment: (a) growth in nutrient broth; (b) growth on nutrient agar; (c) greenish-black colonies with metallic sheen on EMB agar; (d) lactose-fermenting pink colonies on MacConkey agar; (e) yellow colonies on Brilliant Green agar; (f) Gram-negative rod-shaped cells (100x).

### 3.2 Sugar fermentation and biochemical properties

All isolates fermented dextrose, lactose, sucrose, maltose and mannitol with the production of acid and gas, indicated by a colour change from red to yellow and gas bubbles in the inverted Durham’s tubes. Biochemically, all isolates were catalase-positive, indole-positive and methyl red-positive, but Voges–Proskauer-negative and Simmons citrate-negative. On TSI agar, all isolates produced an acidic slant and acidic butt (yellow slant/yellow butt) with gas production, consistent with the standard biochemical profile of E. coli (Table 4).

**Figure 2.**
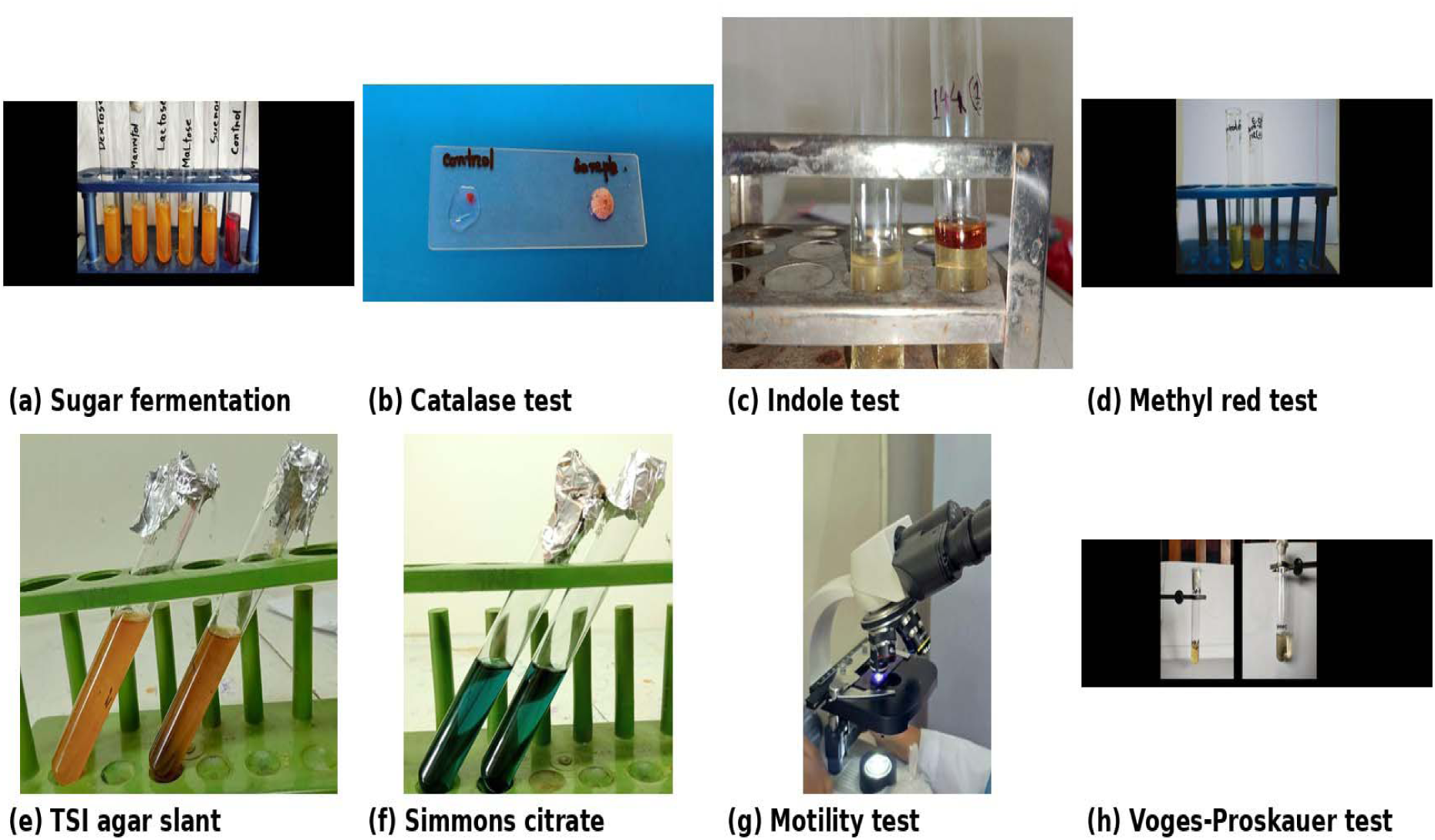
Biochemical characterisation of E. coli isolated from the poultry farm environment: (a) fermentation of five basic sugars with acid and gas production; (b) positive catalase test; (c) positive indole test; (d) positive methyl red test; (e) acid slant/acid butt with gas on TSI agar; (f) negative Simmons citrate test; (g) positive motility (hanging-drop) test; (h) negative Voges-Proskauer test.

**Table 4.**
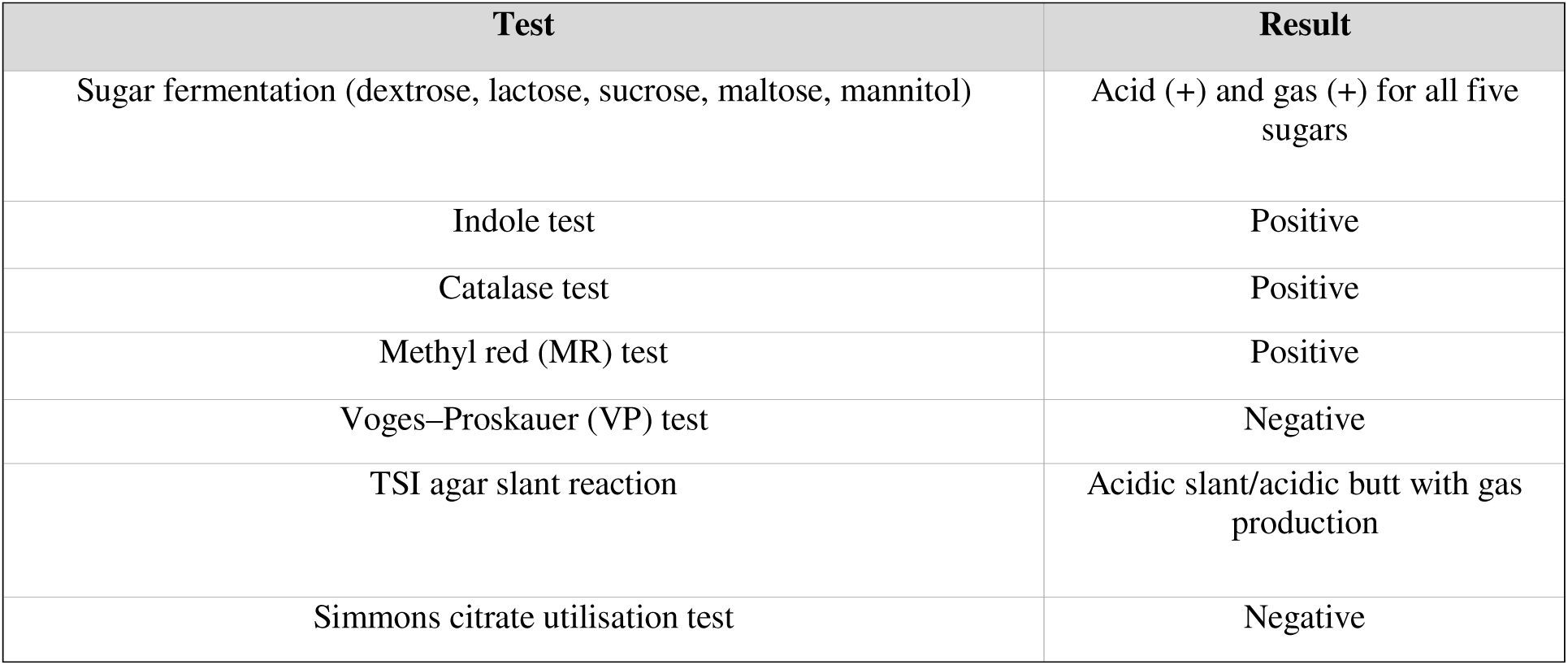
Biochemical profile of the isolated E. coli.

| Test | Result |
| --- | --- |
| Sugar fermentation (dextrose, lactose, sucrose, maltose, mannitol) | Acid (+) and gas (+) for all five sugars |
| Indole test | Positive |
| Catalase test | Positive |
| Methyl red (MR) test | Positive |
| Voges–Proskauer (VP) test | Negative |
| TSI agar slant reaction | Acidic slant/acidic butt with gas production |
| Simmons citrate utilisation test | Negative |

### 3.3 Overall and source-wise prevalence

Out of 60 environmental samples examined, 30 were culturally, morphologically and biochemically confirmed as E. coli, giving an overall prevalence of 50%. Source-wise, prevalence was highest in bird-dropping samples (75%, 15/20), followed by water samples (45%, 9/20) and soil samples (30%, 6/20) (Table 5).

**Figure 3.**
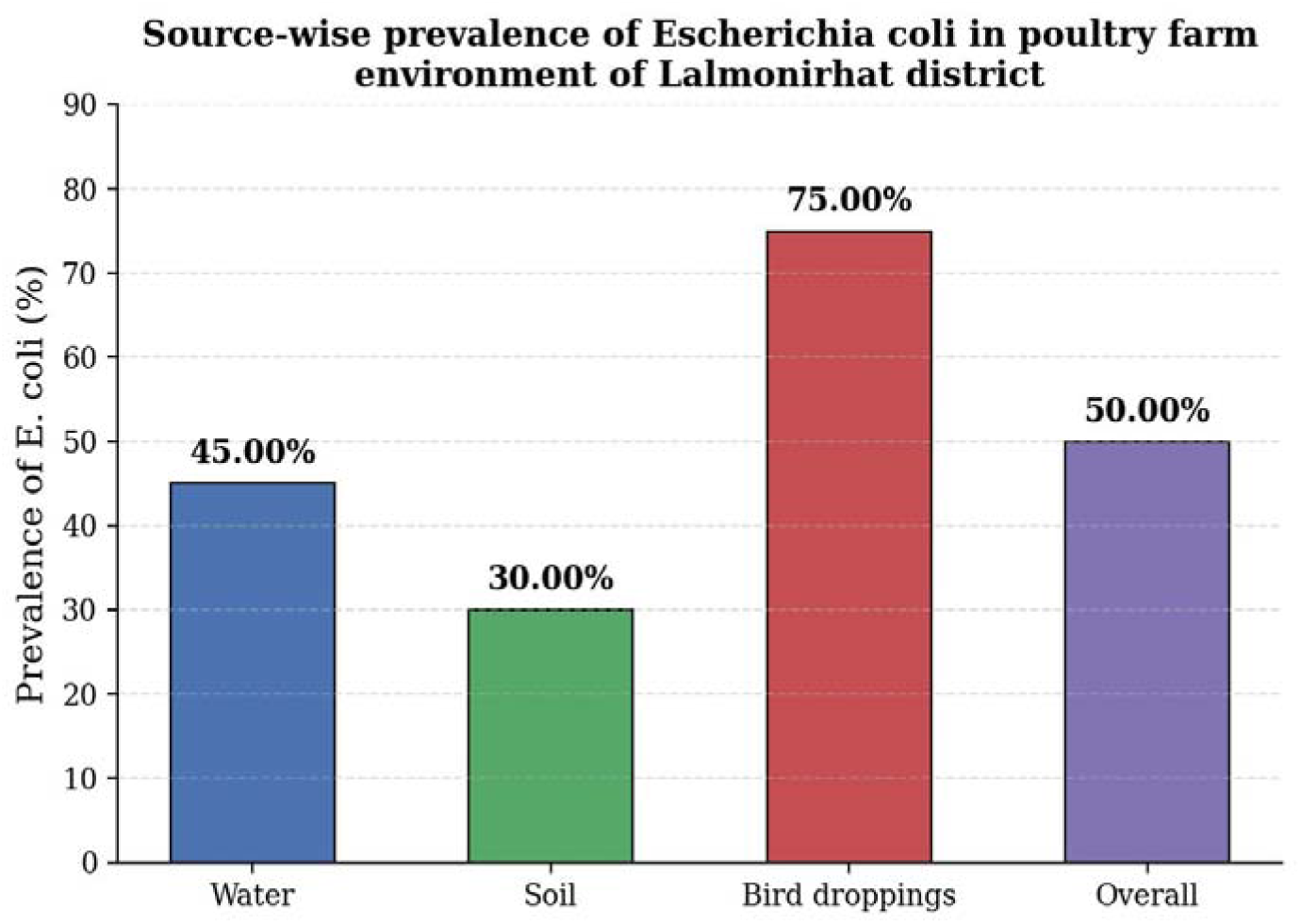
Source-wise prevalence of E. coli in water, soil and bird-dropping samples collected from poultry farms in Lalmonirhat district.

**Table 5.** Overall prevalence of E. coli in environmental samples from poultry farms in Lalmonirhat district.

| Sample source | No. of samples tested | No. positive for *E. coli* (%) |
| --- | --- | --- |
| Water | 20 | 9 (45.00%) |
| Soil | 20 | 6 (30.00%) |
| Bird droppings | 20 | 15 (75.00%) |
| Total | 60 | 30 (50.00%) |

### 3.4 Area-wise (upazila) prevalence

Prevalence of E. coli varied across the five upazilas of Lalmonirhat district, being highest in Hatibandha (66.67%, 8/12) and Sadar (58.33%, 7/12), intermediate in Kaliganj (50%, 6/12) and Aditmari (41.67%, 5/12), and lowest in Patgram (33.33%, 4/12) (Table 6).

**Figure 4.**
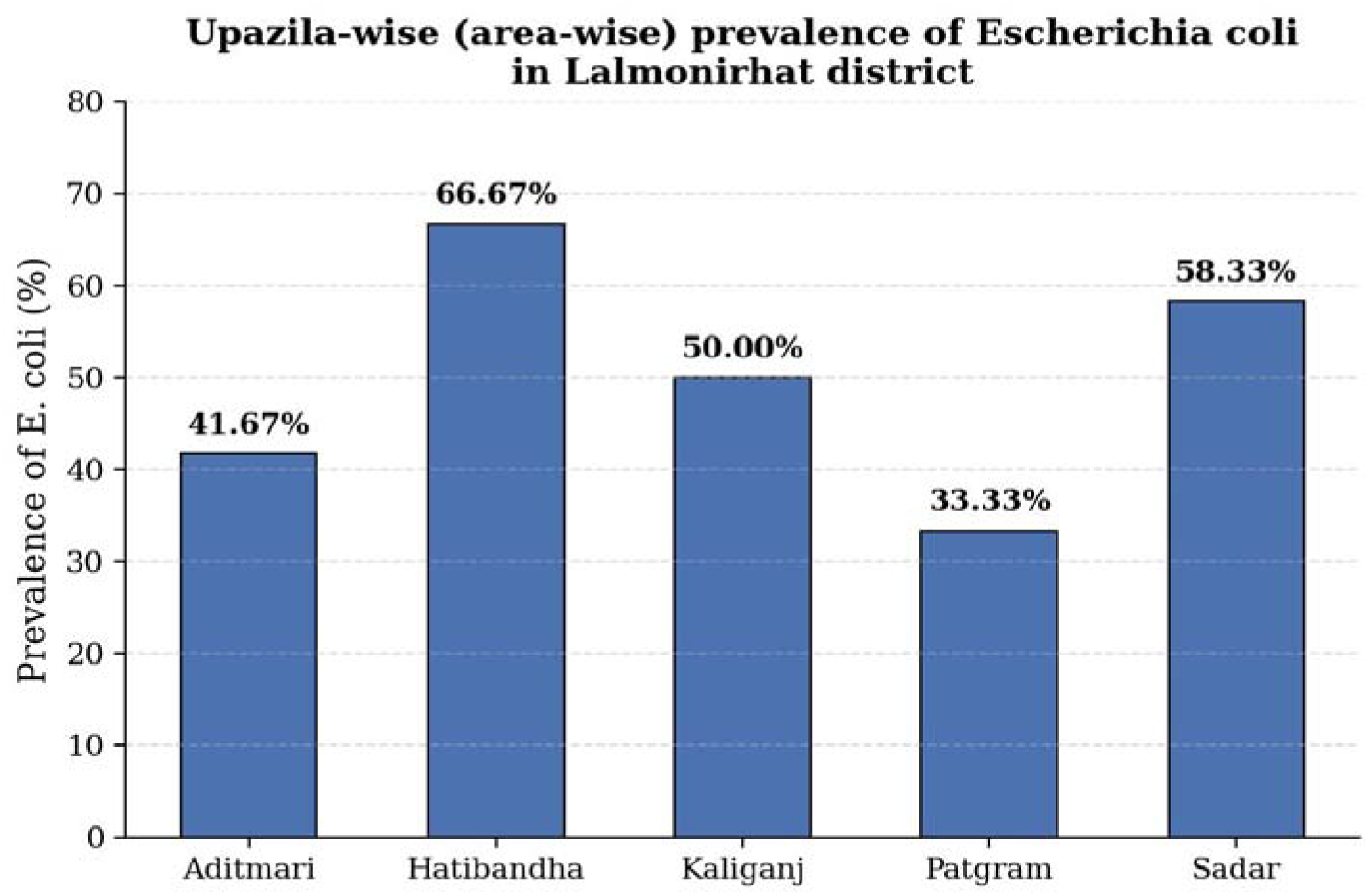
Upazila-wise (area-wise) prevalence of E. coli in the poultry farm environment of Lalmonirhat district.

**Table 6.** Area-wise (upazila-wise) prevalence of E. coli in Lalmonirhat district.

| Upazila | No. of samples tested | No. positive for *E. coli* (%) |
| --- | --- | --- |
| Aditmari | 12 | 5 (41.67%) |
| Hatibandha | 12 | 8 (66.67%) |
| Kaliganj | 12 | 6 (50.00%) |
| Patgram | 12 | 4 (33.33%) |
| Sadar | 12 | 7 (58.33%) |

### 3.5 Antibiogram profile

The 30 confirmed E. coli isolates displayed variable resistance to the seven antibiotics tested. Resistance was highest against oxytetracycline (66.67%) and ciprofloxacin (60%), followed by doxycycline (30%), levofloxacin (16.67%), amoxicillin (13.33%), neomycin (13.33%) and erythromycin (10%). Correspondingly, sensitivity was highest against erythromycin (63.33%), followed by neomycin (56.67%), amoxicillin (56.67%), levofloxacin (50%), ciprofloxacin (30%), oxytetracycline (23.33%) and doxycycline (20%). Intermediate sensitivity ranged from 10% (oxytetracycline and ciprofloxacin) to 50% (doxycycline) (Table 7).

**Figure 5.**
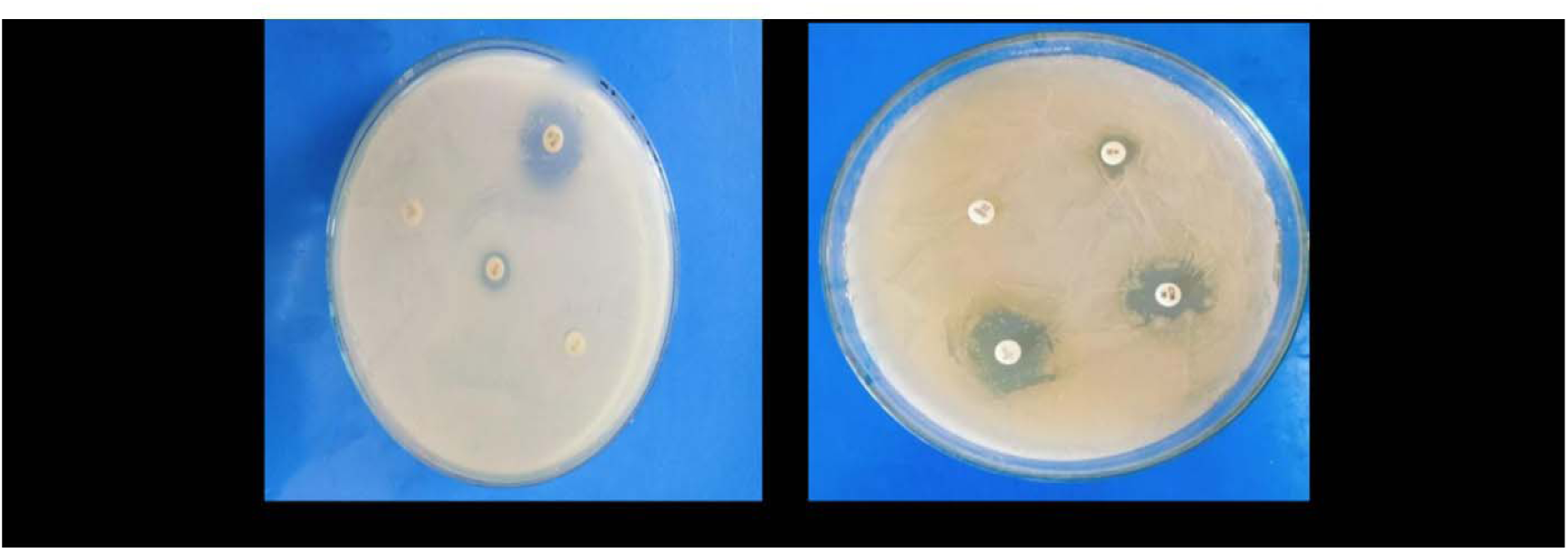
Zones of inhibition produced by E. coli isolates on Mueller-Hinton agar during antibiotic sensitivity testing, showing resistance to some agents and susceptibility/intermediate susceptibility to others.

**Figure 6.**
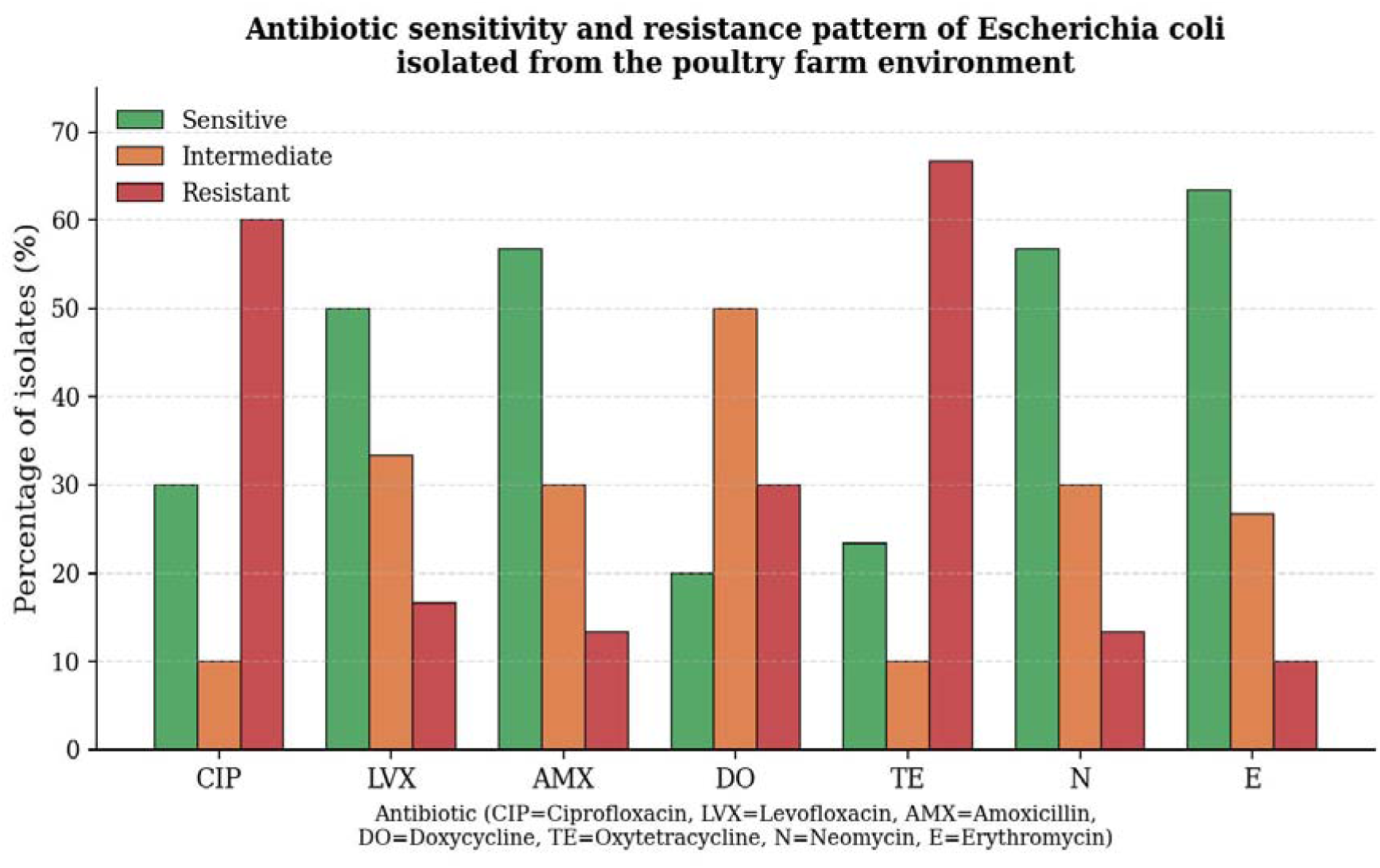
Antibiotic sensitivity and resistance pattern of E. coli isolated from the poultry farm environment (CIP = Ciprofloxacin, LVX = Levofloxacin, AMX = Amoxicillin, DO = Doxycycline, TE = Oxytetracycline, N = Neomycin, E = Erythromycin).

**Table 7.** Antibiotic sensitivity and resistance pattern of isolated E. coli.

| Antibiotic | Sensitive (%) | Intermediate (%) | Resistant (%) |
| --- | --- | --- | --- |
| Ciprofloxacin | 9 (30.00%) | 3 (10.00%) | 18 (60.00%) |
| Levofloxacin | 15 (50.00%) | 10 (33.33%) | 5 (16.67%) |
| Amoxicillin | 17 (56.67%) | 9 (30.00%) | 4 (13.33%) |
| Doxycycline | 6 (20.00%) | 15 (50.00%) | 9 (30.00%) |
| Oxytetracycline | 7 (23.33%) | 3 (10.00%) | 20 (66.67%) |
| Neomycin | 17 (56.67%) | 9 (30.00%) | 4 (13.33%) |
| Erythromycin | 19 (63.33%) | 8 (26.67%) | 3 (10.00%) |

### 3.6 Molecular detection of E. coli by PCR

Of the 30 phenotypically confirmed E. coli isolates, 10 were randomly selected and subjected to PCR targeting the uidA gene. All 10 isolates (100%) produced the expected 486 bp amplicon upon agarose gel electrophoresis and UV visualisation, confirming their molecular identity as E. coli and demonstrating complete concordance between conventional phenotypic identification and uidA-based molecular confirmation (Table 8).

**Figure 7.**
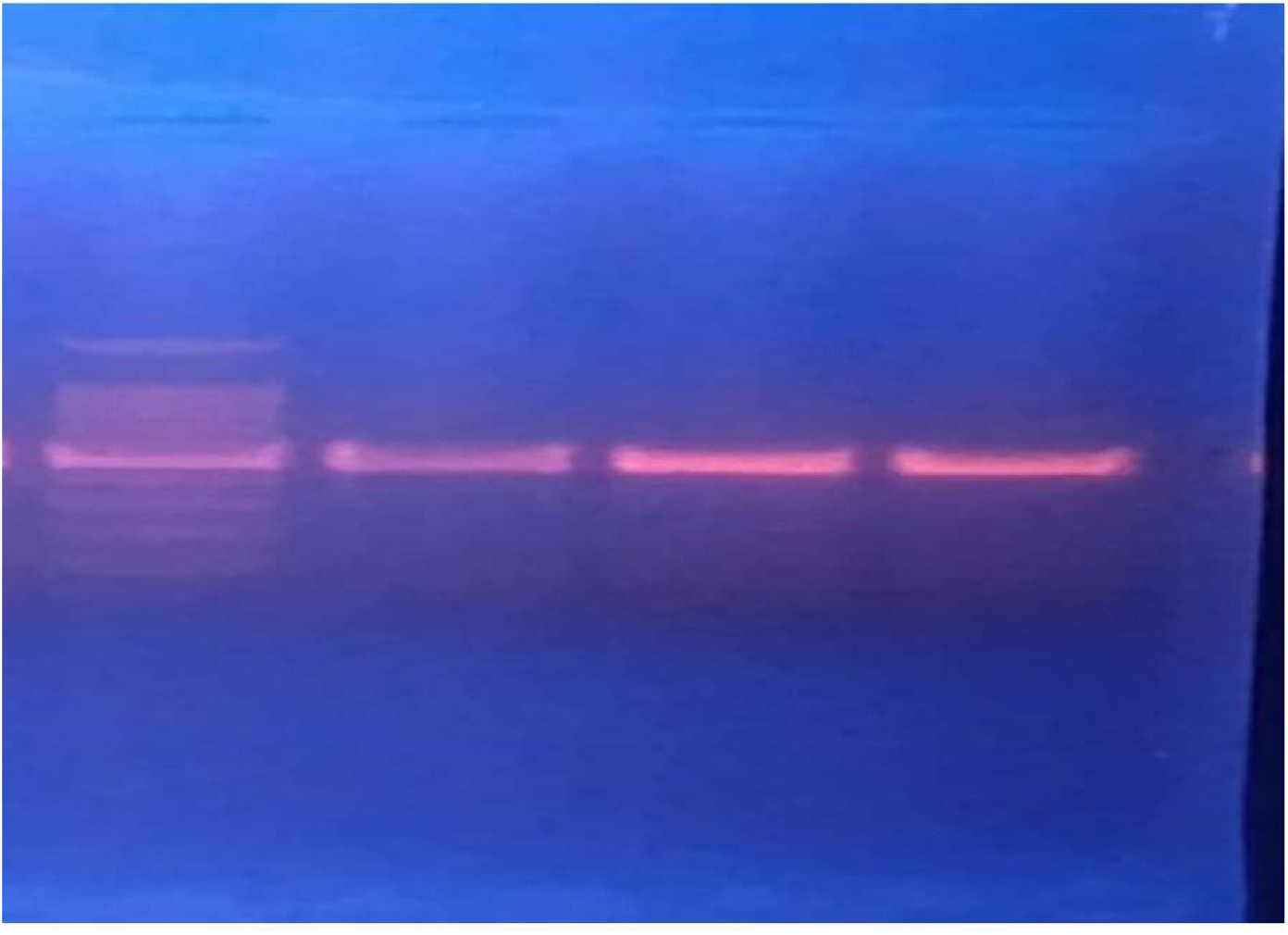
Agarose gel (1.5%) electrophoresis of PCR products showing amplification of the species-specific uidA gene (486 bp) in E. coli isolates from the poultry farm environment.

**Table 8.** Molecular characterisation of E. coli by PCR.

| No. of samples | No. culture-positive | No. of isolates tested by PCR | No. positive for *uidA* gene | Molecular detection rate (%) |
| --- | --- | --- | --- | --- |
| 60 | 30 | 10 | 10 | 100% |

## 4. DISCUSSION

E. coli isolated in this study were Gram-negative, non-spore-forming rods that produced pink colonies on MacConkey agar and a metallic green sheen on EMB agar, consistent with the classical cultural and morphological description of the species (Dou et al., 2016). The 50% overall prevalence of E. coli recorded in the poultry farm environment of Lalmonirhat district is broadly consistent with reports from other regions, although absolute values vary considerably with sample type, season and farm management (Blaak et al., 2015). Zafar et al. (2024) reported a prevalence of approximately 61.33% among 150 samples collected from broiler farms in Faisalabad, Pakistan, while Ibrahim et al. (2023) documented widespread antimicrobial resistance among fecal and environmental E. coli isolates from conventional broiler and Sonali farms in Bangladesh, with virtually all isolates showing multidrug resistance to three to seven antibiotic classes.

The source-wise prevalence pattern observed here- highest in bird droppings (75%), intermediate in water (45%) and lowest in soil (30%)- mirrors the general trend reported across the literature, in which faecal material consistently harbours the greatest bacterial load (Dahshan et al., 2015). Blaak et al. (2015) recorded ESBL-producing E. coli contamination rates of 81% in rinse/run-off water and 60% in dust within poultry-farm environments in the Netherlands, while lower rates were found in surface water, soil and barn air. Da Costa et al. (2008) similarly reported a substantial presence of antibiotic-resistant E. coli in wastewater and sludge from poultry slaughterhouses. Mahmud et al. (2018) found 83.08% of cloacal swabs from apparently healthy broilers in Mymensingh positive for E. coli, and Laube et al. (2014) reported that 86% of slurry samples from German broiler farms were positive for ESBL/AmpC-producing E. coli, with 28.8% of boot swabs and 7.5% of exhaust-air samples also testing positive. Parvin et al. (2020) found E. coli in 71.3% of retail chicken meat and 79.7% of live-bird-market sewage samples across Bangladesh, while Mamun et al. (2016) recorded 81.67% positivity for shigatoxigenic E. coli among cloacal swabs of broilers in Mymensingh, and Dahshan et al. (2015) reported a prevalence of 79.5% in litter, dropping and water samples from Egyptian broiler farms. Collectively, these findings confirm that poultry-farm environments, and faecal material in particular, act as major reservoirs of E. coli, reflecting variation in biosecurity, hygiene and waste-management practices between farms and regions (Jang et al., 2017).

The antibiogram results of the present study revealed the highest resistance to oxytetracycline (66.67%) and ciprofloxacin (60%), with comparatively lower resistance to doxycycline, levofloxacin, amoxicillin, neomycin and erythromycin, interpreted using standard interpretive criteria (CLSI, 2021). This pattern is consistent with the well-documented overuse of tetracyclines and fluoroquinolones in poultry production across South Asia (Habib et al., 2021). Akond et al. (2009) reported that E. coli isolates from poultry and poultry environments in Bangladesh were resistant to penicillin (88%), ciprofloxacin (82%), erythromycin (68%), ampicillin (64%), tetracycline (58%) and chloramphenicol (20%), broadly corroborating the high resistance to older, widely used antibiotic classes observed in the present study. Abdellah et al. (2013) similarly found very high resistance to amoxicillin-clavulanic acid (80%) and norfloxacin (67.5%) among E. coli isolated from turkey meat in Morocco, while El-Boshy et al. (2013) reported an almost identical resistance ranking. Such convergence across geographically distinct poultry systems reinforces the view that the extensive, often unsupervised, use of antibiotics for growth promotion and disease prevention in poultry production is a principal driver of the resistance patterns documented here (Habib et al., 2021; Economou and Gousia, 2015).

Molecular confirmation by PCR provides a rapid, sensitive and highly specific complement to phenotypic identification (Feng, Weagant and Jinneman, 2020). In the present study, the uidA gene, which encodes β-glucuronidase and is widely regarded as a reliable species-specific marker for E. coli, was successfully amplified in all 10 tested isolates, yielding the expected 486 bp product and confirming complete concordance between cultural/biochemical and molecular identification (Sambrook and Russell, 2001). This finding is consistent with the broader literature emphasising the value of PCR-based detection for the rapid and unambiguous confirmation of E. coli in food and environmental samples (Jang et al., 2017; Ievy et al., 2020). The combination of phenotypic and uidA-based molecular methods, as applied here, therefore represents a robust approach for surveillance of E. coli in poultry-farm environments (Choudhari et al., 2020).

Taken together, these findings highlight the substantial and multidimensional public-health risk posed by multidrug-resistant E. coli circulating within poultry-farm environments (Economou and Gousia, 2015). Poor biosecurity, inadequate waste management, and unhygienic handling practices on farms and in surrounding areas facilitate both horizontal transmission among birds and potential spillover to farm workers, consumers and the wider environment (Hossain et al., 2020). Because raw or undercooked poultry products, contaminated water and soil can all serve as vehicles of transmission, addressing this problem requires a coordinated One Health approach that combines improved farm hygiene, prudent and evidence-based antibiotic use, and continuous microbiological surveillance (Dahal, 2017; World Health Organization, 2020).

## 5. CONCLUSION AND RECOMMENDATIONS

This study demonstrates that Escherichia coli is prevalent in the environment of poultry farms in Lalmonirhat district, Bangladesh, with an overall prevalence of 50% and the highest contamination levels in bird-dropping samples. The isolated strains exhibited pronounced resistance to oxytetracycline and ciprofloxacin, moderate resistance to doxycycline, and comparatively greater sensitivity to erythromycin, neomycin and amoxicillin, indicating a substantial burden of antimicrobial resistance associated with commonly used veterinary antibiotics. Molecular confirmation by uidA-targeted PCR corroborated the phenotypic identification of all tested isolates, underscoring the reliability of combining conventional and molecular approaches for E. coli surveillance.

Given these findings, it is recommended that poultry farms in the study area strengthen biosecurity and hygienic practices during production, handling, transport and waste disposal to limit environmental contamination and horizontal transmission of resistant E. coli. Antibiotic use in poultry should be guided by susceptibility testing rather than empirical or prophylactic administration, and alternative disease-prevention strategies such as vaccination and improved farm management should be promoted to reduce reliance on antimicrobials. Continued surveillance integrating phenotypic and molecular methods, together with greater public awareness of the risks of antibiotic misuse, will be essential to curbing the spread of multidrug-resistant E. coli from poultry farms into the wider environment and food chain.

## Acknowledgement

The authors gratefully acknowledge the Department of Veterinary & Animal Sciences, University of Rajshahi, Bangladesh, for providing laboratory facilities and technical support to conduct this research. The authors also sincerely thank the poultry farm owners of Lalmonirhat district for their cooperation during sample collection.

## Ethical Approval

Ethical approval for this study was obtained from the Institutional Animal, Medical Ethics, Biosafety and Biosecurity Committee (IAMEBBC), University of Rajshahi, Bangladesh, before the commencement of the study. Environmental samples were collected with the permission of the respective poultry farm owners, and no live animals were subjected to invasive procedures during the study.

## Informed Consent (Author Consent)

Permission was obtained from the poultry farm owners before sample collection. All authors have read and approved the final version of the manuscript and agree to its submission for publication.

## Data Availability

The data supporting the findings of this study are available from the corresponding author upon reasonable request. All relevant data generated or analyzed during this study are included in this published article.

## Author Contributions

Anna Purnna Ray: Conceptualization, Investigation, Sample collection, Data curation, Laboratory experiments, Data analysis, Methodology, Statistical analysis, Writing- original draft.

Md. Rimon Bhuiyan: Investigation, Data curation, Data analysis, Laboratory experiments, Literature review, Visualization, Writing- original draft, Writing- review and editing.

Satyajit Roy: Data curation, Validation, Data interpretation.

Hridoy Roy: Laboratory investigation, Validation, Data collection.

Md. Sabbir Hossain: Laboratory investigation, Data curation, Validation.

Md. Rashedul Kabir Mondol: Supervision, Project administration, Review and editing, Final approval of the manuscript.

K.M. Mozaffor Hossain: Conceptualization, Methodology, Supervision, Project administration, Writing-review and editing, Final approval of the manuscript.

## Funding

This research received no specific grant from any funding agency in the public, commercial, or not-for-profit sectors.

## Conflict of Interest

The authors declare that they have no competing interests and no conflict of interest regarding the publication of this manuscript.

## Notes

### Competing Interest Statement

The authors have declared no competing interest.

